# Wetting and localization of chromatin condensates by surface-enriched proteins

**DOI:** 10.1101/2025.06.11.658974

**Authors:** Arghya Majee, Jens-Uwe Sommer

## Abstract

We put forward a novel mechanism for the positioning of chromatin within the cell nucleus that does not rely on polymer adsorption to the nuclear lamina. Using a field-theoretic model of a chromatin condensate formed through polymer–protein co-condensation, we show that surface enrichment of chromatin-binding proteins can localize the condensate at the nuclear lamina even when chromatin itself is repelled from the surface. The competition between the tendency of the protein component to wet the surface and the conformational restrictions of the polymer near the impenetrable boundary produces a nonmonotonic chromatin density profile with a maximum at a finite distance from the lamina, consistent with live-cell experiments [Sci. Adv. **7**, eabf6251 (2021)]. We further demonstrate that a non-binding protein competing for surface association can regulate the position of the condensate, providing an alternative pathway by which compositional changes of the lamina can induce chromatin repositioning. A phase diagram of the surface-localization transition reveals how surface affinity, protein availability, and polymer–protein interaction strength control the switch between peripheral and central chromatin configurations. This wetting-based mechanism contrasts with direct polymer adsorption and provides a general framework for understanding how multicomponent condensates interact with responsive biological surfaces.

## I. INTRODUCTION

Polymeric macromolecules like DNA, RNA, and proteins are fundamental to virtually all biological processes. The physical principles governing conformations, phase behaviors and dynamics of these polymers are thus inherently intertwined with key mechanisms in biological systems [1, 2]. One such crucial process is liquid-liquid phase separation (LLPS), a phenomenon that underpins the formation of biomolecular condensates essential for compartmentalizing biochemical reactions and other life functions within cells [3, 4]. These condensates establish distinct microenvironments that not only promote specific cellular processes but also minimize potential interference between concurrently occurring reactions, reduce noise, attract and sequester specific enzymes, and play a fundamental role in the organization of the genome [5].

In eukaryotic cells, the genome is organized into macro-molecular structures primarily composed of DNA and proteins (histones), forming chromatin fibers that can be considered as large, flexible polymers [6]. One mechanism by which condensed chromatin structures – such as heterochromatin, the genetically silenced part of the eukaryotic genome [7] – can form is polymer-assisted condensation (PAC) [8, 9], where flexible polymers act as scaffolds that promote condensation of interacting proteins that would otherwise remain dispersed [10]. While the involvement of polymer phase separation in chromatin organization and its non-uniform distribution across the nuclear volume is now well established, the precise spatiotemporal organization of the genome, along with the factors that govern it, remain poorly understood [11, 12].

Chromatin positioning at the nuclear periphery is often described in terms of lamina-associated domains (LADs), in which specific genomic regions are preferentially associated with the nuclear lamina through protein-mediated interactions [13], with constitutive heterochromatin playing a particularly important role [11, 14]. In such frameworks, peripheral localization arises from direct binding or tethering of chromatin to lamina-associated components. However, it remains unclear whether direct tethering of DNA alone can account for the large-scale relocalization of chromatin observed under varying lamina conditions while preserving the shape and internal organization of chromatin.

Recent *in vivo* experiments in Ref. [15] offer three-dimensional, nuclear-scale insights into chromatin organization in the muscles of live, intact Drosophila larvae. As shown in the left panel of Figure 1(a), under normal conditions, chromatin is preferentially localized at the nuclear periphery, avoiding the central regions of the nucleus. Remarkably, the individual chromosomes remain well separated and do not spread out to form a homogeneous layer along the inner nuclear membrane. However, when the concentration of Lamin A/C, a key protein associated with the nuclear lamina, is increased, chromatin undergoes a structural rearrangement, adopting a condensed conformation at the center of the nuclear volume [right panel of Figure 1(a)]. These observations suggest that the positioning of chromosomes at the nuclear lamina cannot be easily explained by adsorption or desorption of the chromatin polymer. Regardless of the solvent environment, polymer adsorption theory predicts a strong enrichment of polymer density at the surface, with a continuous decay of monomer density toward the center of the nucleus. Consequently, the shape of the adsorbed polymer would resemble a flat “pancake” [16, 17]. Despite this general conclusion from polymer physics, first attempts at explaining these experimental observations focused on invoking adsorption of chromatin polymer to the inner nuclear membrane. In coarse-grained computer simulations, Bajpai *et al*. [18] demonstrated that the adsorption of polymers in poor solvents onto the interior of a sphere can lead to the formation of a chromatin-like layer at the lamina. Upon reducing the adsorption affinity, this state transitions into a droplet-like, unbound polymer conformation, and thus the simulated snapshots qualitatively resemble the behavior of the real system. However, the confinement length scale in these simulations is comparable to the extent of the adsorbed polymer, so that the available surface area limits the lateral spreading of chromatin and can produce an artificial saturation effect. This geometrical constraint can make the adsorbed polymer extend into the interior, qualitatively resembling the experimentally observed peripheral chromosome conformations. In the biological system, by contrast, the nuclear dimensions are on the order of micrometers, and the chromosomes – eight in the case of Drosophila – are distributed along the inner nuclear envelope while retaining an extension of several micrometers perpendicular to the membrane. Such chromosome-scale extension is difficult to reconcile with a conventional polymer adsorption layer, whose thickness is expected to remain microscopic compared with the nuclear dimensions. These observations imply that the problem of chromatin localization at the nuclear periphery – and the experimentally observed chromosome repositioning in response to changes in lamina composition – remains unresolved, suggesting that a mechanism fundamentally different from simple polymer adsorption is at play.

**FIG. 1.**
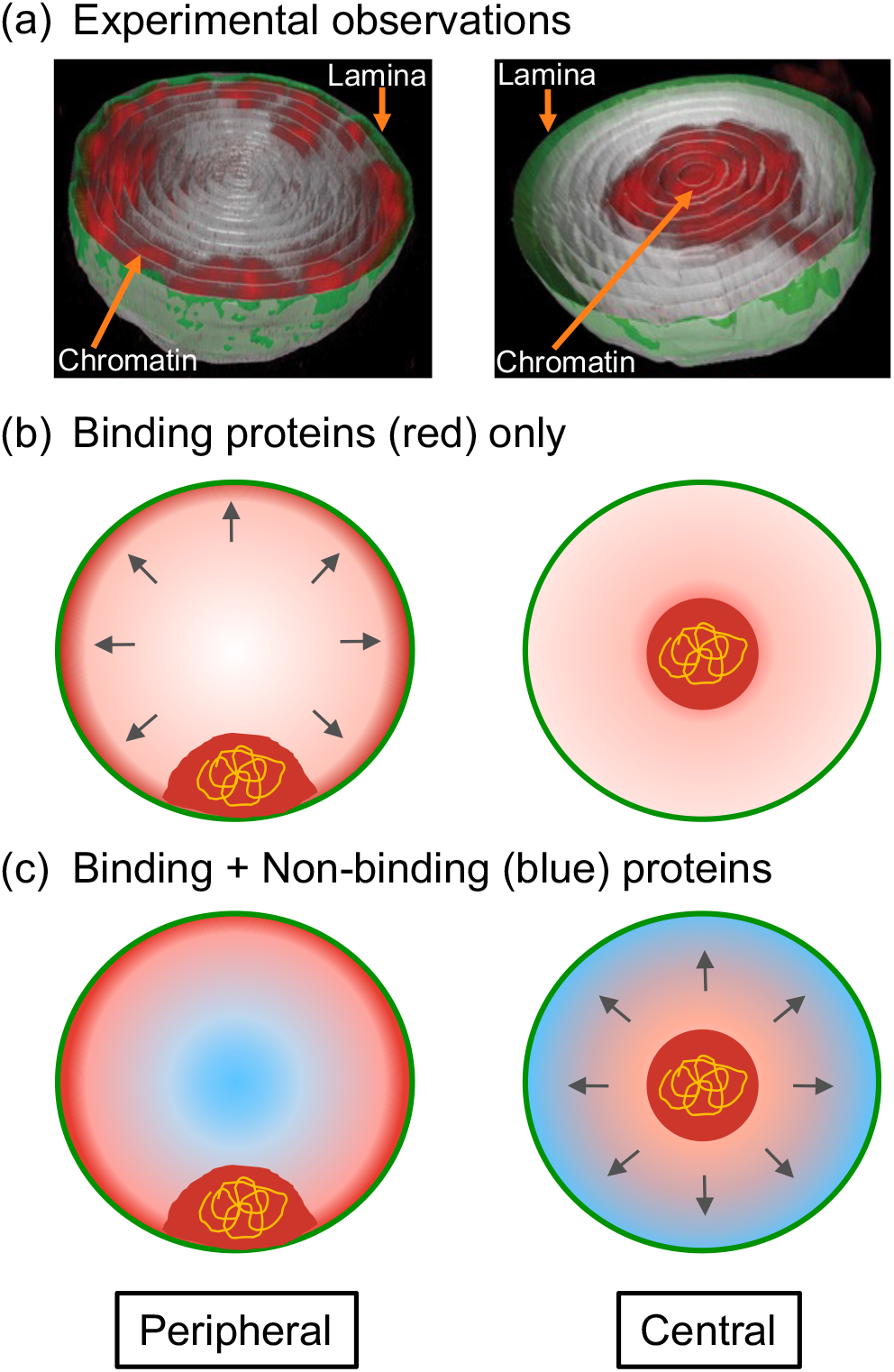
Mechanisms driving chromatin positional changes. (a) Experimental observation of chromatin (red) repositioning in live, intact Drosophila larva (adapted from Ref. [15]): from peripheral localization (left) to centralized configuration (right) upon lamin C upregulation in the nuclear envelope (green). Orange arrows indicate chromatin and lamina regions for clarity. Both panels show nuclear cross-sections. (b) Theoretical model of the phenomenon in panel (a), using a system consisting of a polymer (yellow) with binding proteins (red). As the left panel shows, when these proteins also exhibit affinity for the nuclear lamina (green), a thin protein-rich layer (indicated by the arrows) forms just beneath the lamina, which subsequently attracts the chromatin (a condensate formed by the polymer and binding proteins) towards the lamina. In the absence of this protein–lamina affinity (right panel), no such layer forms, and the chromatin remains centralized. (c) Similar chromatin repositioning as in panel (b) within a model system containing additional non-binding components (blue). When this non-binding component outcompetes binding proteins for lamina affinity (right), they form a new peripheral layer (indicated by arrows) that displaces the binding protein layer, causing the chromatin to localize toward the center of the nucleus. Darker shades denote higher local densities of the depicted components.

In this work, we propose that chromatin positioning at the nuclear periphery is controlled by the surface enrichment of chromatin-binding proteins, rather than by direct adsorption of the polymer to the lamina. Our theory is grounded in the experimental observation that constitutive heterochromatin is preferentially localized at the nuclear lamina [11], with the heterochromatin protein HP1 dynamically responding to changes in lamina composition [19]. While the interaction between HP1 and the lamina involves multiple components – including Lamin A/C [20–23] and the Lamin B receptor (LBR) [24] – notably, HP1 levels are modulated by Lamin C expression, with overexpression of Lamin C leading to a redistribution of HP1 away from the nuclear periphery [19]. These observations motivate our central hypothesis that the association of chromatin-binding proteins such as HP1 with the lamina, for example through interactions with Lamin A/C proteins, can control chromatin localization at the nuclear periphery.

In our framework, chromatin itself is repelled by the lamina, in contrast to adsorption-based models [18, 25] that invoke direct polymer–surface attraction. Peripheral localization instead arises from a wetting phenomenon in which chromatin-binding proteins, which co-condense with chromatin through the polymer-assisted condensation pathway [10], become enriched at the lamina. The resulting protein-rich surface layer stabilizes the chromatin condensate near the lamina even though the polymer itself is excluded from the surface. The competition between protein wetting and the conformational restrictions of the polymer produces a nonmonotonic chromatin density profile with a maximum at a finite distance from the lamina, consistent with the experimental observations [15].

A second difference from collapsed-polymer adsorption models concerns the solvent conditions of chromatin. Such descriptions typically invoke poor-solvent conditions or effective self-attraction to produce polymer collapse, together with an attractive polymer–surface interaction [15, 25]. By contrast, the present mechanism does not require poor-solvent conditions for bare chromatin: condensation is mediated by the chromatin–protein interaction, while the polymer itself remains repelled from the surface. We also note that thermal solvent conditions can be incorporated into the PAC formalism and do not qualitatively alter the transition scenario [10]. This broader applicability is relevant because several established models of chromatin organization represent chromatin as a self-avoiding polymer, with compaction mediated by molecular binders [26–29].

The present surface-localization problem is distinct from our earlier studies of polymer-assisted condensation [10, 30–32], which addressed the formation, internal organization, mechanical response, and biological consequences of polymer-associated condensates, but did not consider their localization at an external surface. Here, the new element is such a surface: it is repulsive to the polymer while favoring enrichment of the binding protein. This asymmetry gives rise to surface-mediated wetting and localization phenomena that are absent from the problems considered in our previous work. We focus on the resulting large-scale redistribution of chromatin between peripheral and central nuclear regions, which is distinct from the internal microphase-like organization of chromatin observed in eukaryotic cells [33, 34]. These forms of chromatin heterogeneity occur on different length scales and may coexist with additional active and structural mechanisms. Our model is therefore intended as a minimal physical description of the interaction of chromatin condensates with the inner nuclear membrane, with particular relevance to constitutive heterochromatin [11, 14].

We explore two pathways by which changes in lamina composition can lead to chromatin repositioning, as illustrated in Figure 1. As shown in panel (b), under normal conditions, chromatin-binding proteins (HP1 in the case of heterochromatin) are enriched at the nuclear periphery – for example, through interactions with Lamin A/C and/or LBR. However, alterations in the composition of the nuclear lamina (such as an increase in Lamin A/C or an associated change in LBR content [35]) can lead to reduced surface enrichment of binding proteins and, eventually, to the relocation of chromatin toward the nuclear center.

Alternatively, this positional shift could be mediated by a *non-binding* molecular component, which does not interact directly with the polymer but instead responds to changes in the nuclear lamina. As illustrated in panel (c), modifications in lamina composition can affect the adsorption behavior of this non-binding component, thereby indirectly altering the local concentration of chromatin-binding proteins at the lamina. A reduction in the surface enrichment of the binding protein – driven by altered adsorption preferences of the non-binding component – could then result in a reorganization of chromatin within the nucleus.

While the mechanism in panel (b) is directly motivated by the experimentally observed response of HP1 to changes in lamina composition [19], panel (c) represents a more general regulatory mechanism based on competitive surface association. We do not assign the non-binding component to a specific molecular species, since such direct competition with HP1 at the lamina has, to our knowledge, not been established.

In this study, we show that surface enrichment of the binding protein can stabilize a chromatin condensate near the lamina even though the polymer itself is repelled from the surface. The resulting competition between protein wetting and polymer conformational restrictions produces a nonmonotonic chromatin density profile with a maximum at a finite distance from the lamina. We characterize the transition between peripheral and central localization and identify the surface affinity and availability of the binding protein, as well as the polymer–protein interaction strength, as key control parameters of this transition. Finally, we emphasize that while our study is motivated by the problem of chromatin organization, it addresses a broader class of wetting and surface-localization phenomena in multicomponent polymer systems, in which different components can have distinct affinities for the surface.

## II. THEORY

We consider chromatin as a coarse-grained polymer system, where the polymer represents the chromatin fiber composed of nucleosome units within a field-theoretic description, i.e. using the density (volume fractions) of the various components as the thermodynamic variables. The local volume fraction of the polymer component (chromatin) is denoted by *ϕ*_*p*_(*z*). Within the ground-state dominance approximation (GSDA), it is convenient to introduce an auxiliary field *ψ*(*z*) such that *ϕ*_*p*_(*z*) = *ψ*^2^(*z*), where *ψ* represents the amplitude of the dominant polymer configuration [36, 37]. In this way, the polymer conformational entropy can be expressed in the standard GSDA form. In addition to the polymer, we include protein components suspended in a solvent, representing chromatin-binding proteins and other nuclear constituents. These proteins can either be binding (interacting attractively with the polymer) or non-binding.

We assume that chromatin organization is radially symmetric. To focus on localization relative to the nuclear lamina, we adopt a one-dimensional Cartesian description in the coordinate *z∈* [0, *L*], where *z* = 0 represents the nuclear lamina and *z* = *L* corresponds to the nuclear interior. The boundary at *z* = *L* is not a physical surface but a symmetry plane with a no-flux (mirror) boundary condition. This one-dimensional description can be viewed as a local flat-wall approximation of the radial problem. It is justified by the separation of length scales between chromatin condensates (with characteristic diameter *∼* 0.1–1 *µ*m) [6, 9, 38–41] and the nucleus (diameter *∼* 5–10 *µ*m) [42], such that curvature effects are small near the lamina where localization is determined. When the condensate is not bound to the lamina, it relocates toward the nuclear interior due to spherical symmetry; in this regime its position is not determined by interaction with a surface, and the boundary at *z* = *L* serves only to represent this symmetric interior region rather than a physical interface.

With these assumptions the free energy *F* per surface unit is given by

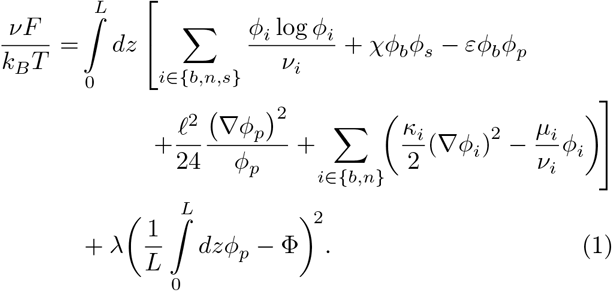

Here, *ϕ*_*b*_(*z*), *ϕ*_*n*_(*z*) and *ϕ*_*s*_(*z*) are the volume fraction profiles for the *b*inding, the *n*on-binding proteins, and the *s*olvent, respectively. The quantities *ν*_*i*_ represent the molecular volumes of the different species in units of *ν*, and control their translational entropy. In particular, larger values of *ν*_*i*_ correspond to reduced mixing entropy per unit volume, as appropriate for coarse-grained macromolecular components. Conservation of volume fractions:

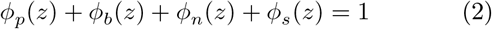

at each spatial location, enables the reformulation of the free energy *F* in Equation (1) as functional of only three variables: *ϕ*_*p*_, *ϕ*_*b*_, and *ϕ*_*n*_. The first term in Equation (1) represents the entropy of mixing of the mobile components, generalized to account for different molecular volume ratios *ν*_*i*_, which reflect the coarse-grained size of each species relative to the solvent reference volume. The subsequent terms correspond to Flory-type interactions: *χ* describes the interaction between binding proteins and solvent, while *ε* captures the interaction between the polymer and the binding proteins. Both quantities are defined in units of thermal energy *k*_*B*_ *T* . The polymer conformational entropy is described within the GSDA framework by the Lifshitz term involving a characteristic length *ℓ* which corresponds to the Kuhn length of the polymer [36, 37, 43, 44]. In this description, the polymer is treated as a connected chain rather than as independent monomers and therefore does not contribute a local ideal mixing entropy term that applies to the small molecular components. The parameter *κ*_*i*_ in Equation (1) is associated with interfacial tensions and the corresponding terms characterize the energy contributions arising from the spatial gradients of constituents in the system [45]. For the sake of simplicity, we have neglected cross-coupling of gradients and higher-order terms involving spatial derivatives.

The protein components are treated in a grand canonical framework, where their concentrations are controlled through chemical potentials *µ*_*i*_. While, at the level of the entire cell, protein copy numbers are regulated, the present description corresponds to an equilibrium coarse-grained model in which the nucleoplasm acts as a reservoir for the proteins. In this sense, the chemical potentials *µ*_*i*_ represent ambient protein concentrations outside the condensate. Phase separation of the proteins alone occurs only at specific coexistence values *µ*_*i*_ = *µ*_*i*,coex_, and in the present work we consider the regime *µ*_*b*_ *< µ*_*b*,coex_, such that the homogeneous dilute phase is the only stable bulk state in the absence of the polymer, see Fig. 2. This ensures that condensation arises through polymer-assisted condensation (PAC) rather than spontaneous protein phase separation. In this mechanism, the presence of the polymer locally modifies the thermodynamic conditions due to the coupling term *− εϕ*_*b*_*ϕ*_*p*_ which represents a local chemical potential trap of depth *εϕ*_*p*_, and induces condensation only within the spatial extent of the polymer conformations (radius of gyration). As a result, condensate formation is spatially confined and does not occur uniformly throughout the nucleoplasm. At the same time, the size of the condensed region is naturally limited by the extent of the polymer, rather than being set by bulk phase separation. We note that condensation of a given component (the binding protein HP1 in our case) occurs only in the presence of a second component (here chromatin). This is a fundamental feature of biomolecular condensates.

**FIG. 2.**
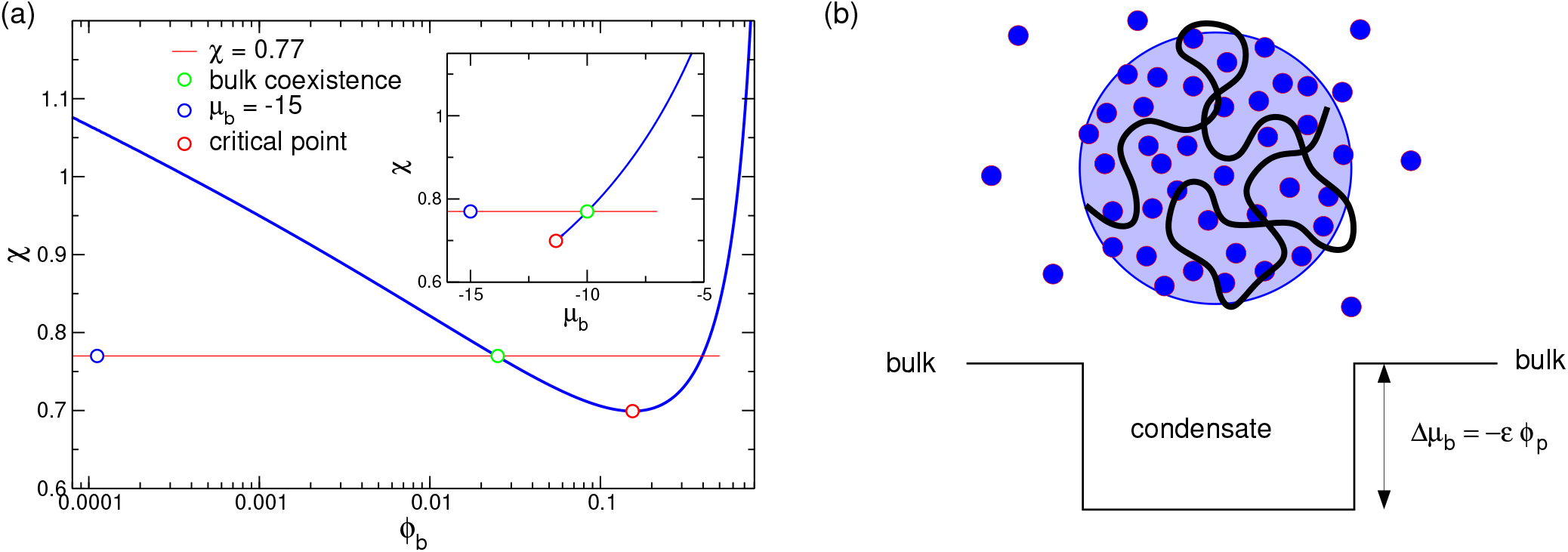
Bulk phase diagram. (a) Phase diagram of the binding protein according to Flory–Huggins theory for a molecular volume ratio *ν*_*b*_ = 30 relative to the solvent. The blue circle indicates the selected bulk state value, while the green circle denotes the volume fraction of the dilute phase at coexistence. (b) Sketch of the physical mechanism of PAC. Polymer conformations create a chemical potential trap for binding proteins. When the local enrichment of proteins within the polymer’s radius of gyration drives the system beyond the coexistence threshold [green circle in (a)], phase separation is induced and a co-condensate with the polymer is formed. Outside the PAC droplet, the proteins remain in the dilute phase [blue circle in (a)]. Thus, interaction with the polymer extends the two-phase region to much lower protein concentrations, but only within the polymer’s radius of gyration. The driving force for droplet formation is therefore weak, nonspecific interactions between the binding proteins, rather than strong, specific bridging interactions.

The monomer volume fraction averaged over the total volume of the system

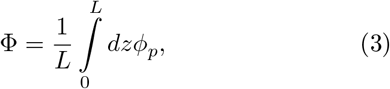

is conserved and *λ* serves as the Lagrange multiplier enforcing this constraint in Equation (1). Thus, Φis the volume fraction of one chromatin polymer averaged over the total volume of the nucleus and should not be considered as the local chromatin density such as in the interior of heterochromatin. Another central part of our polymer physics model is the definition of the appropriate boundary conditions. In contrast to previous work on chromatin adsorption we assume that chromatin is not adsorbed at the lamina and thus the monomer concentration vanishes at *z* = 0 [16]. The other components representing the proteins satisfy Neumann (natural) boundary conditions at the surfaces, which reflect their affinity for the surface [46]:

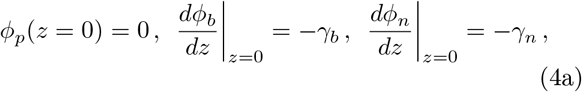

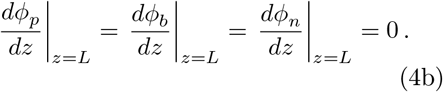

The parameters *γ*_*b*_ and *γ*_*n*_ describe the affinity of the respective protein species to the lamina and control their enrichment or depletion at the boundary. These parameters represent effective surface affinities that incorporate the different molecular contributions to the localization of the respective components at the lamina, rather than affinities of specific molecular pairs. In this coarse-grained framework, these parameters represent effective surface interactions [46, 47] rather than microscopic binding energies, and are chosen to span regimes from depletion to strong adsorption. The inner wall of the nucleus is repulsive to the chromatin polymer, whereas the binding protein can become enriched at the boundary. At the same time, the binding protein forms a dense phase only through its attractive coupling (of strength *ε*) to the polymer, since its bulk concentration remains below the coexistence concentration. The resulting wetting problem is therefore intrinsically multicomponent: the component attracted toward the surface is not itself capable of forming the bulk condensate, while the polymer required for condensation is excluded from the surface. Surface localization consequently emerges from the coupled redistribution of the two components rather than from direct adsorption of the polymer.

The equilibrium density profiles are obtained by minimizing the free energy functional in Equation (1) with respect to the fields *ϕ*_*p*_(*z*), *ϕ*_*b*_(*z*), and *ϕ*_*n*_(*z*), subject to the constraint in Equation (2) and the boundary conditions in Equations (4). This is equivalent to solving the associated Euler–Lagrange equations *δF/δϕ*_*i*_ = 0, with *i∈{p, b, n}*, which determine the equilibrium density profiles. In practice, this minimization is carried out using a gradient-descent algorithm on a discretized domain (see Methods for details). We note that for the homogeneous system the analytical calculation of the corresponding phase diagrams is possible, and we refer here to previous work [10, 30, 31].

## III. RESULTS AND DISCUSSIONS

First, we consider a system that contains only polymer and binding proteins in a solvent, i.e., non-binding proteins are not present. The molecular volumes are chosen as *ν*_*s*_ = *ν* = 1 for the solvent, corresponding to a coarse–graining volume of 1 nm^3^, i.e., about 30–35 water molecules. This choice reflects the fact that the nucleoplasm is a crowded environment containing a wide range of molecular species, such that treating the solvent as uncorrelated pure water molecules would overestimate its translational entropy. The reference volume *ν* should therefore be interpreted as an effective coarse-graining volume that accounts for this compositional complexity. Taking into account that HP1 has a molecular volume corresponding to roughly 10^3^ water molecules, this coarse-graining yields *ν*_*b*_ = 30 for the binding proteins.

The polymer is characterized by an average monomer volume fraction Φ= 0.08 and *ℓ* = 10 nm, which corresponds to typical extension of a nucleosome. According to Eq. (3), Φrepresents a spatially averaged chromatin volume fraction over the entire system, including both chromatin-rich and chromatin-depleted regions. This value appears consistent with reported chromatin volume fractions in eukaryotic cells [48]. We can estimate the volume of a nucleosome with a diameter of about 10 nm and a height of about 5 nm [49] with approximately 400 nm^3^ which results in about 2 nm^3^ per base pair. For the genome of Drosophila we can take approximately 3 *×* 10^8^ bp which results in a total dry volume of about *µ*m^3^. Given a typical nuclear volume of about 80 *µ*m^3^ for the nucleus yields a chromatin volume fraction of roughly 0.01 which may provide a lower estimate for Φ. We note that experimental measurements typically probe local densities within subnuclear volumes and reveal strong spatial heterogeneity. Accordingly, the local chromatin density within the condensate in our model is significantly higher than the global average as will be reported below.

The Flory parameter, which characterizes the interaction between the proteins and the solvent, is set to *χ* = 0.77. For the specific case *ν*_*b*_ = 30 the critical point for the binding protein is located at 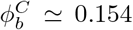 and *χ*^*C*^*≃* 0.7, see Fig. 2. The coexistence chemical potential *µ*_coex_ is obtained by solving the standard conditions of equal chemical potential and osmotic pressure between coexisting phases, corresponding to the common-tangent construction of the free energy density. For the chosen parameters, this yields *µ*_*b*,coex_ *≈−* 10. We choose the chemical potential of the binding proteins as *µ*_*b*_ = *−*15, which lies below the coexistence value and corresponds to volume fraction of about *ϕ*_*b*_ *≃* 0.0001. Using the coarse-grained units described above, the bulk concentration of the binding protein is therefore set to about 5.5 *µ*M. For a typical nuclear volume of 80 *µ*m^3^, this corresponds to approximately 2.7 *×* 10^5^ binding proteins. The choice of the interaction parameter is more difficult to validate against experimental observations. However, *in vitro* studies report a saturation threshold of several hundreds of *µ*M for HP1*α*,[8], which leads to a ratio between the bulk value and the saturation value on the order of 100. Taking into account that the bulk concentration of proteins should be lower in coexistence with the condensate due to the finite size of the nuclear volume as compared to the averaged value, our ansatz for a ratio of about 200 appears to be close to the estimated range. We note that increasing the parameter *χ* in our free energy model (thus reducing the saturation density) shifts the system deeper into the PAC region and thus strengthens the observed condensation and wetting scenario. The same holds for increasing the bulk concentration at a given interaction parameter, see Fig. 2. We conclude that the choice of our model parameters is consistent with the available experimental data [50].

With the given parameters, the homogeneous dilute phase is the only stable bulk state of the protein system, and phase separation does not occur spontaneously in the absence of the polymer. However, due to the attractive interaction between the polymer and the binding proteins, characterized by *ε* = 1.8 in Equation (1), the local thermodynamic conditions within the spatial extent of the polymer are modified. This drives the system locally into the two-phase region and induces condensation via polymer-assisted condensation (PAC) [10]. A useful way to interpret this mechanism is to view the polymer as creating a local environment that effectively shifts the protein chemical potential toward coexistence. As a result, condensation occurs only in regions where the polymer is present. This leads to spatially confined condensates whose size is naturally limited by the extent of the polymer, rather than being determined by bulk phase separation. For the numerical examples presented below, we use a gradient penalty parameter *κ*_*b*_ = *κ*_*n*_ = 60 nm^2^, which sets the interfacial width. This choice is consistent with previous studies [51–53] and yields physically reasonable interface thicknesses. All energies are expressed in units of *k*_*B*_ *T*.

Figure 3 illustrates the spatial variations of the volume fractions of different components for various values of the parameter *γ*_*b*_ which describes the affinity of the binding protein to the lamina located at *z* = 0, see Equation (4a). Larger values of *γ*_*b*_ lead to stronger accumulation of proteins at the surface, whereas smaller values correspond to weaker or vanishing adsorption (*γ*_*b*_ = 0). This behavior is clearly shown in Figure 3: as the value of *γ*_*b*_ is gradually reduced from panel (a) to (c), the binding protein concentration *ϕ*_*b*_(0) at the wall diminishes. Notably, the location of the chromatin condensate – marked by increased volume fractions of both the polymer and proteins – also shifts in response to changes in *γ*_*b*_. As panel (a) suggests, the condensate prefers to stay close to the surface when the proteins strongly accumulate at the lamina, forming a thin layer of elevated protein density at *z* = 0. For smaller values of *γ*_*b*_ [panel (b)], the surface enrichment is weaker, and the condensate moves away from the lamina. Notably, the condensate is located in the nuclear interior already at finite surface affinity of the binding protein which indicates the subtle interplay between polymer entropy which favors a delocalization and wetting tendency of the protein. The proximity of the condensate to the surface can be attributed to the attractive interaction between the surface-enriched proteins and the polymer that forms the condensate. However, as seen in panel (b), mere protein adsorption at the surface is insufficient to stabilize the condensate near the lamina: only when the attraction is sufficiently strong [panel (a)] it can overcome the loss of polymer conformational entropy near the boundary. For weaker surface affinity, the reduced protein enrichment no longer stabilizes the condensate at the lamina. We note that the spatial extent of the chromatin-rich condensate in Fig. 3 reflects the amount of polymer considered in the calculation. As discussed below, increasing the polymer amount primarily increases the extent of the condensate while leaving its local composition essentially unchanged. The approximately 100 nm scale observed here should therefore not be interpreted as an upper length scale imposed by the PAC mechanism.

**FIG. 3.**
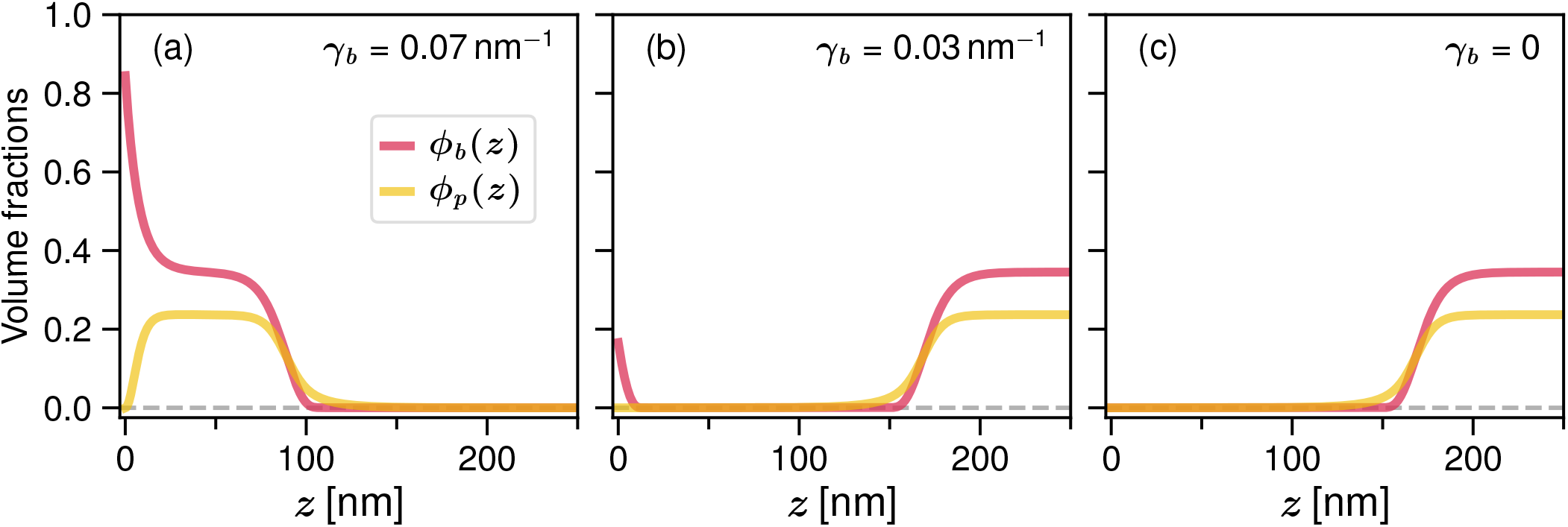
Positional Shifts of Condensates Induced by Protein Adsorption on a Surface. Location of a condensate, formed through attractive interaction between proteins and a polymer, as function of the affinity of the protein to adsorb at a surface characterized by the parameter *γ*_*b*_ defined in Equation (4a). The volume fraction profiles for the monomers, *ϕ*_*p*_, and for the binding protein, *ϕ*_*b*_ are displayed. (a) When the proteins exhibit a strong attraction to the surface (*γ*_*b*_ = 0.07), a thin protein layer forms adjacent to the surface, effectively drawing the condensate to remain in close proximity. (b) As the protein affinity for the surface diminishes (*γ*_*b*_ = 0.03), a layer still forms at the surface, but the concentration of proteins within the layer remains relatively low. Consequently, the attractive force exerted by the layer is insufficient to retain the condensate. Due to the symmetry of the system, the condensate migrates toward the mirror plane at *z* = *L* = 250 nm. (c) In the absence of adsorption (*γ*_*b*_ = 0), proteins do not accumulate at the surface, eliminating the attractive interaction needed to hold the condensate near the lamina. Consequently, the condensate stabilizes at the center of the system, marked by the mirror plane. For the plots, we have used molecular volumes *ν* = 1 and *ν*_*b*_ = 30, monomer volume fraction Φ= 0.08, Flory parameter *χ* = 0.77, a negative chemical potential for the binding protein in the bulk *µ*_*b*_ = *−* 15, a protein-polymer binding strength of *ε* = 1.8, a surface parameter *κ*_*b*_ = 60 nm^2^, and *ℓ* = 10 nm. It is noteworthy that proteins with a chemical potential below the coexistence value, as is the case considered here, can still undergo phase separation due to the polymer acting as a potential trap [10].

We emphasize the distinction between the present wetting mechanism and the classical surface adsorption of a collapsed polymer. In the latter case, an excess layer of adsorbed monomers is always formed at the surface, leading to a monotonically decaying density profile, with a significant fraction of the monomers residing within a thin adsorption layer [17]. This conventional monotonic behavior is not shown here. In contrast, all profiles displayed in Figures 3(a)–(c) exhibit non-monotonic variation of the monomer volume fraction consistent with experiments that report chromosome localization near the lamina without full adsorption. In particular, live-imaging experiments by Amiad–Pavlov et al. [15] show peripheral chromatin organization with enrichment near the nuclear lamina, while maintaining depletion toward the nuclear interior rather than a simple monotonic decay expected for adsorption (see their Figures 1, 3, and 4), as well as a transition from peripheral to more centrally localized chromatin upon Lamin overexpression (their Figure 5). Similarly, in our model, the chromatin density increases from zero at the lamina to a maximum at a finite distance from the surface due to protein-mediated attraction and then decreases toward the nuclear center. The depletion of chromatin directly at the lamina is imposed through the standard Dirichlet boundary condition *ϕ*_*p*_(0) = 0. However, this condition only fixes the depletion directly at the wall and does not determine whether the chromatin condensate localizes near the surface or in the nuclear interior. The finite-distance maximum in the chromatin density therefore arises naturally from the interplay between polymer exclusion from the surface and its attraction to the surface-enriched binding proteins, without requiring an additional, specifically constructed polymer– surface “bonding” boundary condition, as introduced by Adame-Arana and Safran [25]. More importantly, the surface interaction in the present model acts on a component distinct from the polymer. The surface enriches the chromatin-binding protein, while chromatin itself remains excluded. Consequently, wetting and localization are controlled by the coupled composition of the condensate and can be regulated through the surface affinity or availability of the binding protein, rather than through a polymer–surface interaction alone. Furthermore, as we will demonstrate below, this provides a straightforward mechanism for controlling surface localization through the introduction of a competing protein that interacts with the surface but does not bind to the condensate polymer. To elaborate on this more general competitive mechanism, we introduce an additional component that does not bind to the polymer but can compete with the binding protein for surface association. This non-binding protein component has a significantly lower chemical potential of *µ*_*n*_ = *−* 45, corresponding to a vanishingly small bulk concentration, such that it appears only through surface enrichment. The non-binding protein is assigned the same molecular volume as the binding protein, *ν*_*n*_ = 30, and we use identical gradient parameters *κ*_*b*_ = *κ*_*n*_ = 60 nm^2^. All other parameters are kept identical to the previous case unless otherwise specified.

**FIG. 4.**
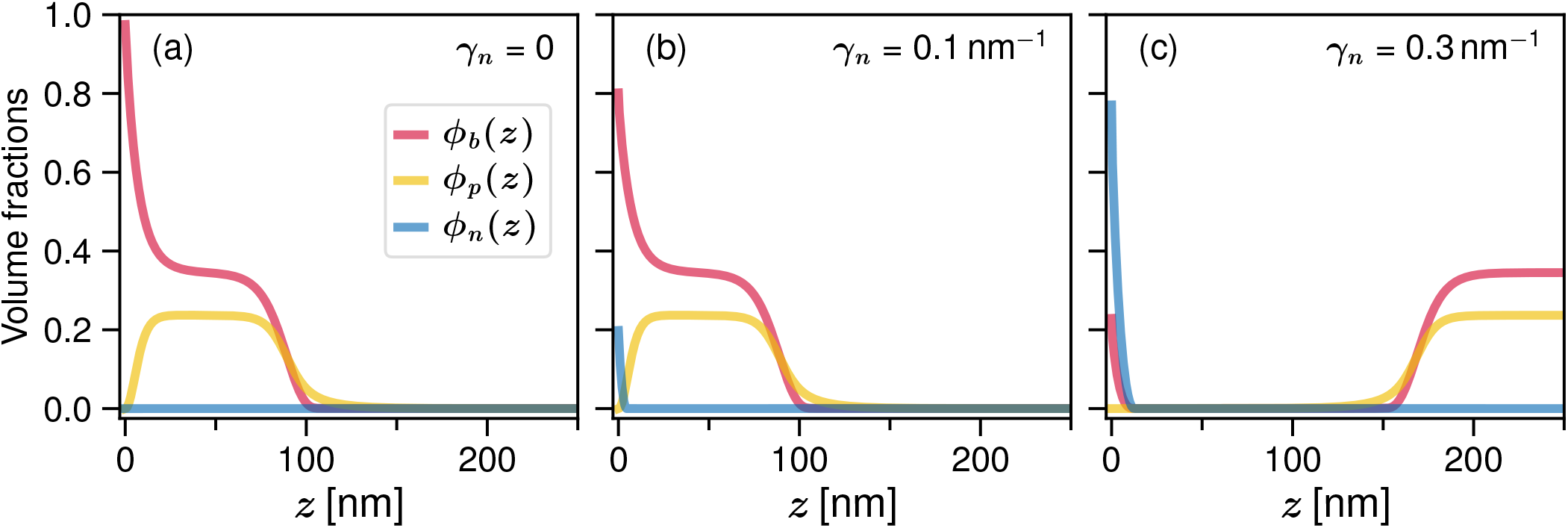
Positional Shifts of the Condensate Induced by interplay of different protein adsorptions onto bounding surface. Location of a condensate, formed due to attractive interaction between proteins and a polymer, in response to varying affinities of a protein that does not directly interact with the condensate components, but adsorbs at the nuclear boundary. The adsorption affinity of the proteins are characterized by the parameters *γ*_*b*_ and *γ*_*n*_, as defined in Equation (4a). For all the plots a fixed value of *γ*_*b*_ = 0.1 nm^*−*1^ is used and *γ*_*n*_ is varied relative to that. (a) For *γ*_*n*_ = 0 *< γ*_*b*_, the non-binding protein does not adsorb at the surface, and the binding proteins are strongly enriched at the lamina, forming a thin layer that attracts and stabilizes the condensate near the surface. (b) For *γ*_*n*_ = *γ*_*b*_ = 0.1, both protein species have comparable affinity for the surface. The non-binding protein begins to compete with the binding protein for adsorption, reducing the surface concentration of the latter. The condensate remains near the surface, but the stabilizing interaction is weakened. (c) For *γ*_*n*_ = 0.3 *> γ*_*b*_, the non-binding protein dominates the surface adsorption, significantly reducing the surface enrichment of the binding protein. As a result, the attractive interaction required to stabilize the condensate at the lamina is lost, and the condensate relocates toward the nuclear interior, marked by the mirror plane at *z* = *L* = 250 nm. For the plots, we have used a Flory parameter *χ* = 0.77, a negative chemical potential *µ*_*n*_ = *−* 45, and surface parameters *κ*_*b*_ = *κ*_*n*_ = 60 nm^2^. All other parameters are the same as in Figure 3.

**FIG. 5.**
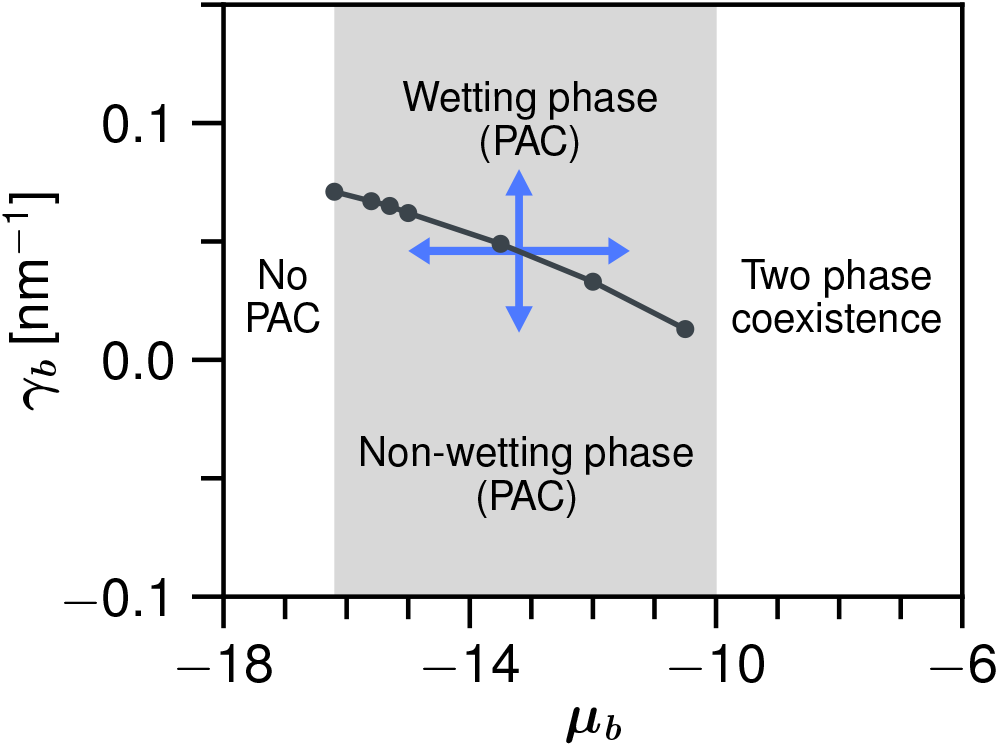
Surface phase diagram. Phase diagram including the surface states. The gray-shaded region identifies the parameter space where droplet formation is driven by PAC. As indicated by the blue arrows, within this PAC region, the droplet undergoes a transition from a central (non-wetting) to a peripheral (wetting) configuration when the interaction parameter *γ*_*b*_ increases, or, when the chemical potential *µ*_*b*_ is raised within a specific range of *γ*_*b*_. To the right of the gray-shaded region, droplet formation occurs independently of PAC, while to the left, droplet formation is entirely suppressed. Unless otherwise stated, the parameters are consistent with those in Figure 3.

In Figure 4, we modulate the affinity of the non-binding protein for the surface by adjusting the parameter *γ*_*n*_ while keeping the affinity *γ*_*b*_ of the binding protein constant. This mimics the depletion of the binding protein at the lamina by a regulatory (non-binding) component.

As panel (a) shows (*γ*_*n*_ = 0 *< γ*_*b*_), the non-binding protein does not adsorb to the surface, and the binding protein remains strongly enriched at the lamina. This results in a sufficiently strong attractive interaction to keep the condensate localized near the surface. In the case *γ*_*n*_ = *γ*_*b*_ [panel (b)], both protein species exhibit comparable affinity for the surface. As a result, the non-binding protein begins to compete with the binding protein for adsorption sites, leading to a reduction in the surface concentration of the binding protein. Consequently, the attraction acting on the condensate is weakened. Still, the binding protein is predominantly present at the surface, where it generates a sufficiently strong attraction to keep the condensate near the surface. However, when the affinity of the non-binding protein exceeds that of the binding protein [*γ*_*n*_ *> γ*_*b*_, panel (c)], the non-binding component dominates the surface. This significantly suppresses the surface enrichment of the binding protein and eliminates the attractive interaction required to stabilize the condensate at the lamina. As a result, the condensate shifts toward the nuclear interior (mirror boundary).

An analogous localization threshold therefore exists for the non-binding competing component. At fixed *γ*_*b*_, increasing *γ*_*n*_ eventually produces sufficient competitive surface enrichment of the non-binding component to suppress the enrichment of the chromatin-binding protein and destabilize the peripheral condensate. We have also examined how this threshold depends on the chemical potential *µ*_*n*_ of the competing component. Increasing *µ*_*n*_, corresponding to a greater availability of the competing species, lowers the value of *γ*_*n*_ required to induce the wall-to-center transition. Thus, competitor availability and surface affinity act together: when more competitor is available, a smaller *γ*_*n*_ is required to produce sufficient competitive adsorption at the surface to suppress the enrichment of the chromatin-binding protein.

The transition between a surface-localized and a central condensate can be understood as a free-energy crossover between two alternative configurations. Surface localization is favored by adsorption of the chromatin-binding protein at the lamina and its attractive interaction with the polymer. This energetic gain competes with the cost of placing the polymer close to a surface from which it is excluded, where its available conformations are restricted. Moreover, because the binding protein is attracted to the surface whereas the polymer cannot contact it, the peripheral state involves a local compositional rearrangement of the PAC condensate, producing a protein-rich and polymer-poor region close to the wall. Surface enrichment of the binding protein is therefore an important factor favoring peripheral localization, but its magnitude alone does not determine the transition. The localization threshold is set by the overall free-energy balance between surface adsorption, polymer–protein attraction, polymer conformational entropy, and the near-surface compositional rearrangement.

The chromatin depletion-layer thickness is not an independently prescribed parameter in our model. The boundary condition *ϕ*_*p*_(0) = 0 enforces depletion directly at the lamina, while the spatial extent of the depleted region emerges from minimization of the complete free energy. Its characteristic width is determined by the combined effects of the polymer conformational-gradient term, the protein gradient penalty, the polymer–protein interaction, and the surface interaction. The depletion-layer width can therefore contribute to the free-energy balance associated with surface localization, but it is an emergent property of the equilibrium profiles rather than an independent parameter that fixes the transition.

To further examine the role of protein availability in the transition between surface and central configurations, Fig. 5 presents a phase diagram as a function of the chemical potential *µ*_*b*_ and the surface interaction parameter *γ*_*b*_ of the binding proteins. For simplicity, we consider a system without non-binding proteins.

As observed, the gray-shaded region for *µ*_*b*_ *< µ*_*b*,coex_ corresponds to droplet formation driven by PAC. Within this region, we varied *γ*_*b*_ for each value of *µ*_*b*_ and determined from the equilibrium density profiles the crossover points between central and surface-localized droplets. The solid line with dots in Figure 5 connects these numerically obtained points and thus delineates the boundary between the two configurational states. For *µ*_*b*_ *> µ*_*b*,coex_, the binding proteins can phase separate even in the absence of the polymer, leading to the usual two-phase coexistence between a dense protein-rich phase and a dilute surrounding phase, as obtained from standard mean-field coexistence conditions [10, 30]. For sufficiently negative *µ*_*b*_, no condensation occurs at all, since both the PAC-driven and the conventional droplets vanish. Once the lower chemical-potential limit is reached for a particular *ε*, no droplet formation is observed upon further decreasing *µ*_*b*_.

For fixed *µ*_*b*_, increasing the surface interaction strength *γ*_*b*_ within the PAC region induces a transition from a central (non-wetting) to a peripheral (wetting) condensate. Conversely, at fixed *γ*_*b*_, the transition can also be triggered by increasing *µ*_*b*_. A larger *µ*_*b*_ increases the availability of binding protein, resulting in higher protein concentrations in the PAC condensate and facilitating protein enrichment near the surface. The resulting protein profile is determined self-consistently, since surface adsorption, polymer–protein attraction, and the energetic cost of protein-density gradients all contribute to the free-energy balance. Consequently, a smaller surface affinity can be sufficient to stabilize the peripheral state at larger *µ*_*b*_.

The localization transition also depends on the chromatin–protein interaction strength *ε*. At fixed *γ*_*b*_, increasing *ε* can change the equilibrium configuration from surface-detached to surface-attached. We find that increasing *ε* increases the binding-protein concentration within the PAC condensate, while its polymer density and size remain essentially unchanged. At the same time, larger *ε* directly strengthens the attractive coupling *− εϕ*_*b*_*ϕ*_*p*_ between chromatin and binding protein. Together, these effects shift the free-energy balance in favor of surface localization. In the parameter range examined, increasing *ε* therefore lowers the threshold value of *γ*_*b*_.

We also examined the effect of the mean chromatin volume fraction Φ. Varying Φfrom 0.06 to 0.10 produces no appreciable change in the localization threshold. Changing Φprimarily changes the spatial extent of the PAC condensate, while the polymer and binding-protein volume fractions inside and outside the condensate remain essentially unchanged. Consequently, the local composition of the PAC condensate and its interaction with the surface-enriched protein layer are largely unaffected. We note, however, that in the present single-polymer formulation, increasing Φby increasing the polymer content at fixed system size also corresponds to increasing the length of the single polymer. Chromatin concentration and polymer length are therefore not independent control parameters under this variation. Varying chromatin concentration independently of polymer length, for example by changing the number of polymers at fixed polymer length, would require a multi-chain extension of the model.

Finally, protein size enters the theory through the molecular-volume parameter *ν*_*b*_, which explicitly controls the translational-entropy contribution of the binding protein. Changing *ν*_*b*_ also modifies the equilibrium protein and polymer density profiles and therefore indirectly affects the other free-energy contributions. We examined the localization transition over the range *ν*_*b*_ = 10–60 for several values of *µ*_*b*_. The localization threshold changes only weakly over this range and shows no systematic monotonic dependence on *ν*_*b*_, whereas its dependence on *µ*_*b*_ is substantially stronger. Thus, within the parameter range examined, the localization threshold is relatively insensitive to the protein molecular volume.

## IV. CONCLUSION

In summary, we have introduced a wetting-based mechanism that offers a potential explanation for chromatin positioning within the nucleus. In particular, we show how a chromatin condensate, such as heterochromatin, can be localized near the lamina without requiring either direct adsorption of chromatin to the surface or explicitly poor-solvent conditions for the chromatin polymer. The key ingredient is the asymmetric surface response of the condensate components: chromatin is repelled from the lamina, whereas chromatin-binding proteins can become enriched at the surface. The attraction between the surface-enriched binding proteins and chromatin draws the condensate toward the nuclear periphery, while the repulsive boundary excludes chromatin from the lamina itself. The competition between these effects produces a nonmonotonic chromatin density profile with a maximum at a finite distance from the surface. Importantly, the mechanism remains valid if bare chromatin exhibits some degree of self-attraction, and our conclusions are therefore largely independent of its solvent conditions. This robustness further extends to the choice of coarse-graining parameters: we have verified that using equal molecular volumes for all components, which shifts the system to significantly higher concentration regimes, leads to the same qualitative behavior.

The surface-localization mechanism proposed here is not mutually exclusive and may act in conjunction with LAD-based tethering and other regulatory processes, providing an additional physical route to control chromatin organization within the nucleus. More broadly, chromatin organization is influenced by multiple concurrent physical processes, and polymer phase separation can coexist with active non-equilibrium mechanisms such as loop extrusion and chromatin remodeling [54–58]. We note that the pronounced segregation into chromatin-rich and chromatin-depleted regions described here reflects large-scale organization driven by surface interactions, as observed in the Drosophila muscle-cell nuclei considered here.

Our results reveal several routes for regulating the surface localization of the condensate. Changes in the surface affinity or bulk concentration of the chromatin-binding proteins, as well as in the strength *ε* of their attraction to chromatin, can shift the condensate between peripheral and central configurations. Moreover, regulatory components – proteins that do not bind to the polymer but compete with the binding proteins for surface association – can provide an indirect route for condensate repositioning. These results highlight the importance of multicomponent effects in controlling the interaction of biomolecular condensates with surfaces. While our primary focus here is on chromatin organization, it is worth noting that the implications of our theory may extend beyond this specific context, particularly in light of the widespread presence of biological condensates and the growing interest in how these condensates interact with and adhere to various surfaces [59–66].

The present analysis, based on a one-dimensional mean-field description, captures the essential physics governing chromatin–lamina interactions. Nevertheless, real chromatin condensates exhibit three-dimensional droplet morphologies. Preliminary molecular dynamics simulations of spherical chromatin droplets qualitatively support the trends predicted here. A comprehensive analysis of these 3D dynamics will be presented in an upcoming manuscript.

Extending this framework to multicomponent systems, such as those recently explored in simulations by Tangade et al. [67], could reveal how compositional heterogeneity and competitive interactions shape interfacial organization within the nucleus. Moreover, our findings may inspire new approaches to the controlled adsorption of polymers at substrates in mixed solvent environments.

## ACKNOWLEDGMENTS

The authors acknowledge support from the Deutsche Forschungsgemeinschaft (DFG) under the grant SO 277/25. J.U.S. acknowledges support by the DFG under Germany’s Excellence Strategy-EXC2068-390729961-Cluster of Excellence Physics of Life. The authors also acknowledge the Center for High-Performance Computing (ZIH) Dresden for granting simulation time.

## AUTHOR CONTRIBUTIONS

The analytical model was developed by both authors.

A.M. has performed the numerical calculations. Both authors contributed equally to manuscript writing.

## APPENDIX: METHODS

The free energy in Equation (1) is minimized numerically on a one-dimensional discretized domain *z ∈* [0, *L*] using a gradient descent algorithm. All fields depend only on the coordinate *z*, and the gradient operators in Eq. (1) therefore reduce to derivatives with respect to *z*. The integrals are evaluated using the trapezoidal rule and spatial derivatives are represented within a finite-difference approximation. The implementation is carried out using a custom C++ code.

The minimization is performed with respect to the spatially varying fields *ϕ*_*p*_(*z*), *ϕ*_*b*_(*z*), and, when present, *ϕ*_*n*_(*z*). For numerical convenience, the polymer field is represented by an auxiliary variable *ψ*(*z*) defined through *ϕ*_*p*_(*z*) = *ψ*^2^(*z*), such that the free energy is minimized in terms of *ψ*(*z*), *ϕ*_*b*_(*z*), and *ϕ*_*n*_(*z*). The solvent volume fraction *ϕ*_*s*_(*z*) is determined from the local incompressibility condition,

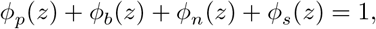

which, in terms of the auxiliary field, reads

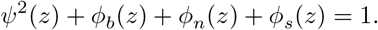

In the two-component case (polymer + binding protein), this reduces to *ψ*^2^(*z*) + *ϕ*_*b*_(*z*) + *ϕ*_*s*_(*z*) = 1. The mirror-symmetry boundary condition at *z* = *L* is implemented as *∂*_*z*_ *ψ* = 0, which implies *∂*_*z*_ *ϕ*_*p*_ = 0. To improve numerical stability and to ensure that all volume fractions remain within their physical bounds, the minimization is carried out in terms of transformed variables. The transformed variables *u, v*, and *w* are functions of *z*; for notational simplicity, this dependence is omitted in the following expressions. The polymer field is parametrized as

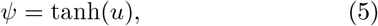

which guarantees 0 *≤ ψ*^2^ *≤*1. The remaining available volume fraction 1 *− ψ*^2^ is then distributed among the molecular species by logistic-type transformations.

For the polymer + binding-protein system, we use one additional variable *v*, such that

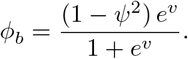

For the polymer + binding-protein + non-binding-protein system, we use two additional variables *v* and *w*, such that

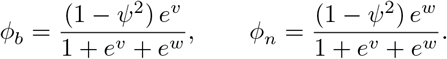

This parametrization guarantees positivity of all components and automatically enforces the local volume-fraction constraint.

The iterative minimization is performed by gradient descent in the transformed variables. Denoting a generic transformed field by *q*, the update step reads

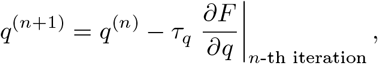

where *τ*_*q*_ is a step size that may differ for the different fields to account for their relaxation scales. In practice, the derivatives are evaluated using all fields at their values at iteration *n*, i.e., (*u*^(*n*)^, *v*^(*n*)^) in the two-component case and (*u*^(*n*)^, *v*^(*n*)^, *w*^(*n*)^) in the three-component case.

The optimization is considered converged when the free energy no longer changes appreciably between successive iterations. In practice, the optimization was continued until the change in free energy between successive iterations became negligibly small. For all reported results, the absolute change in free energy between the last two iterations was typically of order 10^*−*10^ or smaller, corresponding to relative changes of order 10^*−*11^ or below.

To reduce the risk of convergence to metastable states, many randomized initial profiles are tested, and the configuration yielding the lowest free energy is selected. The global constraint on the monomer volume fraction in Eq. (3) is enforced through the penalty term proportional to *λ* in Eq. (1), such that deviations from the prescribed average monomer volume fraction incur a large free-energy cost.

